# Arabidopsis AZG2, an auxin induced putative cytokinin transporter, regulates lateral root emergence

**DOI:** 10.1101/2020.01.31.927970

**Authors:** Tomás M. Tessi, Sabine Brumm, Eva Winklbauer, Benjamin Schumacher, Carlos I. Lescano, Claudio A. González, Dierk Wanke, Verónica G. Maurino, Klaus Harter, Marcelo Desimone

## Abstract

The phytohormones cytokinin (CK) and auxin are key regulators of plant growth and development. During the last decade specialised transport mechanisms turned out to be the key for the control of local and long distance hormone distributions. In contrast to auxin, CK transport is poorly understood. Here we show that *Arabidopsis thaliana* AZG2, a member of the AZG purine transporter family, acts as CK transporter involved in the determination of the root system architecture. The expression of *AtAZG2* is primarily auxin dependent and restricted to a small group of cells surrounding the lateral root primordia. Compared to wild type, mutants carrying loss-of-function alleles of *Atazg2* have higher density of lateral roots, suggesting AZG2 as being part of a regulatory pathway in lateral root emergence. Moreover, *azg2* mutants are partially insensitive to exogenously applied CK, which is consistent with the observation that the CK reporter gene *TCSn_pro_:GFP* showed lower fluorescence signal in the roots of *azg2* mutants compared to those of wild type. These results indicate a defective CK signalling pathway in the region of lateral root primordia. By the integration of AtAZG2 subcellular localization and CK transport capacity data, our results allowed us to propose a local Auxin/CK signalling model for the regulation of lateral root emergence.

## INTRODUCTION

The shape of the plant body results from a fine tuned interaction between genetic programs and environmental conditions. Root development is a great example for this: from an embryonic organ, plants modulate the root architecture in accordance to its requirement for anchorage, nutrients or water uptake. Root branching relies on the capability to secondary develop lateral roots (LR). This process begins with the mitotic activation of founder cells in the pericycle and the subsequent formation of a lateral root primordia (LRP) (De Smet et al., 2007). For lateral root emergence, LRPs have to overcome the mechanical resistance offered by the external cell layers. However, it remains unclear how all these tissues interact with each other during LR organogenesis (Lucas et al., 2013).

Lateral root development is tightly coordinated by different signals with an outstanding role of the morphogenic hormones auxins and cytokinins (CK). Changes in their biosynthesis, transport, perception and crosstalk are necessary events for a proper LR development (Nordstrom et al., 2004; Bishopp et al., 2011; Marhavy et al., 2014).

A crucial event in lateral root initiation is the generation and maintenance of high auxin concentrations in the primed pericycle cells and primordial tip cells (Benkova et al., 2003). The auxin concentrations at cellular level are modulated by cell to cell transport through polar localised transporters (Grones and Friml, 2015). Polarized distribution of these transporters in the plasma membranes of these cells is regulated by a complex signalling network, in which CK plays a crucial function (Laplaze et al., 2007; Marhavy et al., 2014). Auxin transport is also involved in lateral root overlaying tissues (OLT) remodelling which facilitate lateral root emergence. For instance, the AUXIN RESPONSE FACTORs (ARF) 7 and 19 modulate the transcription factor LATERAL ORGAN BOUNDARIES DOMAIN 29 (LBD29) which in turn transcriptionally regulates cell wall remodelling enzymes and transporters such as LAX3 (Swarup et al., 2008, Porco et al., 2016). Consistently, the double *arf7-1/arf19-1* and the single *lbd29* and *lax3* loss-of-function (LOF) mutants show lateral root developmental deficient phenotypes (Okushima et al., 2005; Wilmoth et al., 2005; Swarup et al., 2008; Feng et al., 2012). In contrast to auxin, the role of CK in the OLT remodelling remains unclear. External addition of CK reduces the number of emerged lateral roots in a dose dependent manner whereas overexpression of CK degrading enzymes as well as CK perception deficient mutants show the opposite effect, indicating a potential restrictive function of CK in lateral root emergence (Bertell and Eliasson, 1992, Werner et al., 2010, Riefler et al., 2006). Nevertheless, it is not clear yet if CK restricts the lateral root emergence at the OLT level and, in case, how this process is coordinated with auxin signalling and function.

In contrast to the abundant information on auxin transport, little is known about CK transport. Despite the described processes involving CK mobilization (Samuelson et al., 1992; Takei et al., 2004; Kiba et al., 2013; De Rybel et al., 2014; Ohashi-Ito et al., 2014; Chang et al., 2015), only a few putative cytokinin transporters were reported for a long time (Burkle et al., 2003, Hirose et al., 2005; Sun et al., 2005). However, no strong evidence was provided to link these transporters to physiological processes (Kudo et al., 2010). Lately, the ABC transporter AtABCG14, and the purine transporter PUP14 were described as plasma membrane localized CK transporters involved in long distance transport and in morphogenesis, respectively (Ko et al., 2014; Zhang et al., 2014 Zürcher et al., 2016).

This work is focused on the function of transporters belonging to the AZA-GUANINE RESISTANT (AZG) family. The first member of this transporter family has been described in *Emerichella nidulans,* EnAZGA, as a purine transporter. Some years ago, *Arabidosis thaliana* AZG1 and AZG2, were identified and briefly characterized as adenine-guanine transporters (Mansfield et al., 2009). However, their *in vivo* function was not studied up to date. In this work, we provide evidence that *AZG2* is expressed in a small number of cells surrounding the LRP. Furthermore AZG2 is transporting with high efficiency not only purines but also to *trans*-zeatin (tZ) and other CK, suggesting that the transporter could be involved in CK regulation of lateral root development. Moreover, transcriptional, phenotypic and CK reporter gene studies of *azg2* LOF mutants and plants ectopically expressing AZG2 revealed that the transporter is an inhibitory factor within the auxin/cytokinin hormonal crosstalk network involved in the regulation of lateral root emergence.

## RESULTS

### AZG2 is expressed around lateral root primordia in the overlaying tissues

Transcriptomic studies (Arabidopsis eFP Browser; Winter et al., 2007) and RT-PCR analysis showed that *AZG2* mRNA was mainly present in roots, but also in reproductive tissues (Figure 1A and Supplemental Figures 1C). To analyse the tissue specific expression pattern of *AZG2* in more detail, *AZG2_pro_:GUS* and *AZG2_pro_:GFP* stable transgenic Arabidopsis plants were generated. Strong GUS and GFP signals were detectable in a few cells of the root’s OLT throughout lateral root development (Figures 1B and 1C) but not in reproductive organs like flowers, pollen and seeds (Supplemental Figures 2A to 2D). Nevertheless, weak GUS staining was visible in micropylar endosperm cells of germinating seeds (Supplemental Figures 2E to 2G). To gain more insight on *AZG2* expression in the root, three dimensional reconstructions of cross sections of *AZG_pro_:GFP* expressing root tissue were performed. GFP fluorescence was detected along the primordia at the region where cells became separated (17µm) to allow the emergence of the new lateral root (Figure 1Db). Thereby, reporter activity was observed in cortical and epidermal cells (Figure 1Dc; segmented lines). This distinctive expression pattern in lateral root primordia OLT suggests that AZG2 may play a role in the regulation of lateral root development.

**Figure 1.**
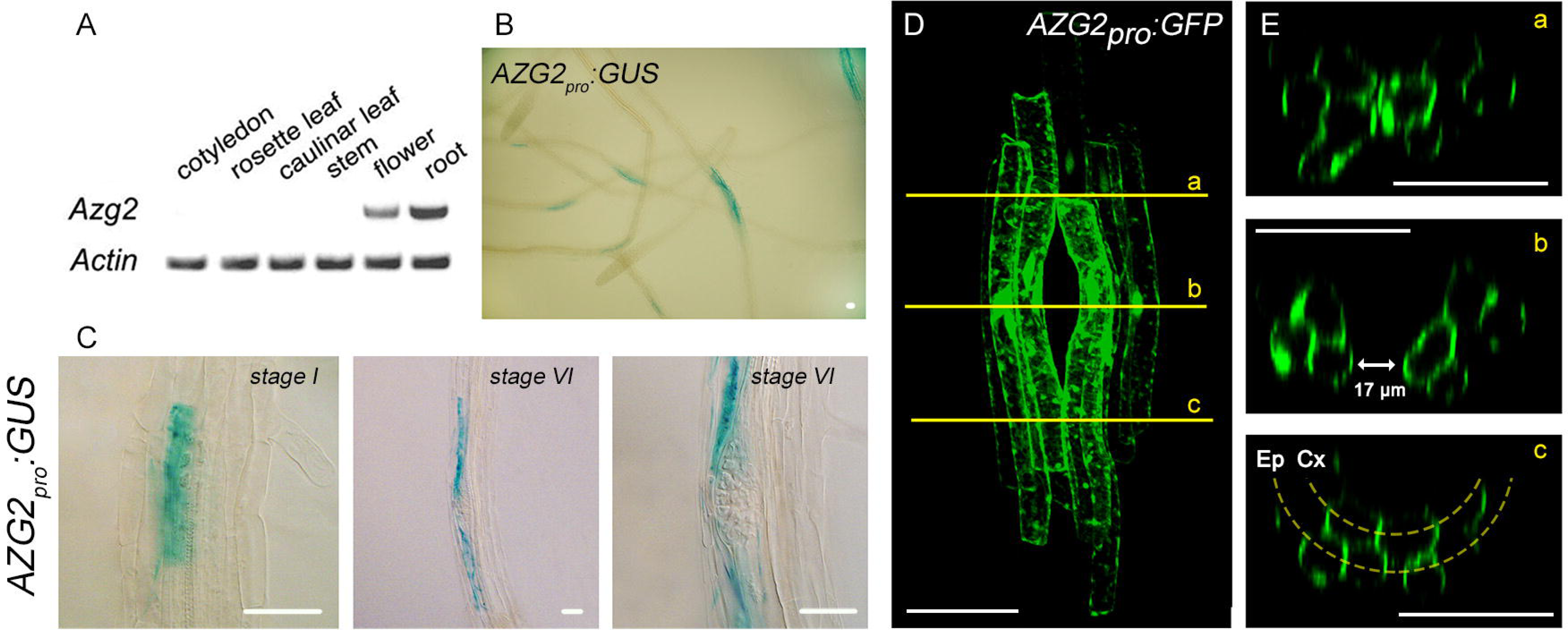
*AtAZG2* is mainly expressed in the overlaying tissues around lateral root primordia. (A) Semi-quantitative RT-PCR analysis of *AtAzg2* expression in various plant tissues with *Actin2* as loading control. AZG2 transcript is detectable in flowers and roots. **(B)** GUS staining of transgenic *AZG2_pro_:GUS* Arabidopsis plants reveals strong AZG2 expression around the lateral root primordia in roots. **(C)** Early (*stage I*) and late (*stage VI*) stages of LRP showing expression of *AZG2_pro_:GUS.* Scale bar, 100µm. **(D)** Three dimensional reconstruction of cells expressing *AZG2_pro_:GFP* around a lateral root primordia, including three different cross sections (a, b and c; yellow lines represents the transverse planes; Ep: Epidermis; Cx: Cortex) . Scale bar, 50µm.

**Figure 2.**
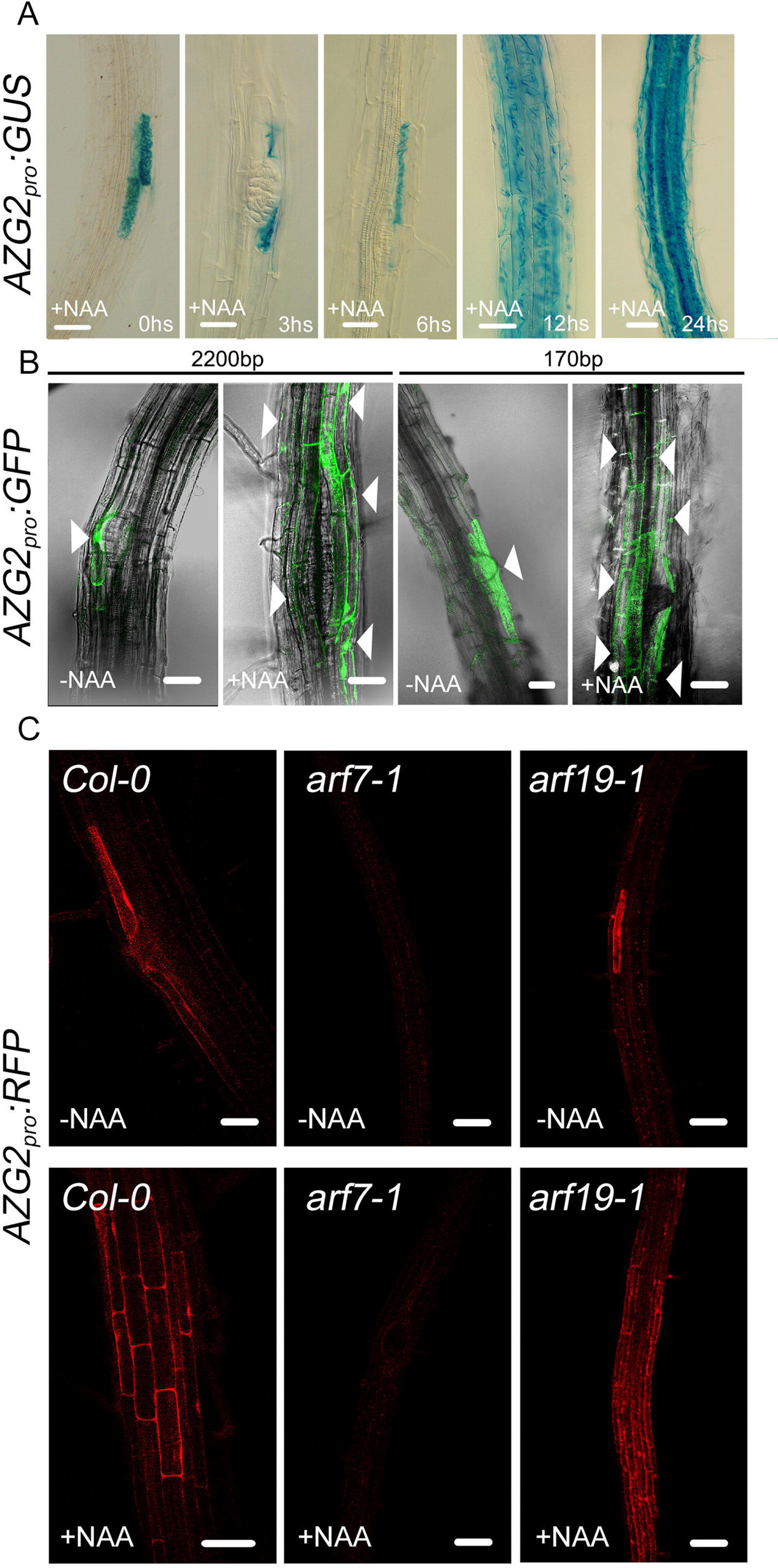
*AtAzg2* expression is auxin inducible (A) 1µM 1-Naphthaleneacetic acid (NAA) treatment of *AZG2_pro_:GUS* transgenic Arabidopsis roots for 0, 3, 6, 12 and 24 hours shows strong expression induction. **(B)** Activity of two *AtAZG2* promoter length (2200 and 170bp) driving GFP expression after a 12 hour treatment with and without NAA treatment. Both promoter lengths show AZG2 expression and induction *via* auxin **(C)** Expression analysis of *AZG2_pro_:RFP* in different genetic backgrounds (*Col-0*, *arf7-1* and *arf19-1*) after a 12 hour treatment with and without NAA. *AtAZG2* is expressed in *Col-0* and *arf19-1* genetic background, but not in *arf7-1*. Scale bars,100 µm.

### Auxin induces AZG2 expression *via* ARF7 transcription factor

As auxin is one of the major hormones involved in the transcriptional regulation of lateral root development genes, it was studied whether auxin has an influence on *AZG2* expression as well. Microarray data analysis revealed that *AZG2* expression was induced 3 hours after of IAA treatment (Supplemental Figures 3A; Winter et al., 2007; Goda et al., 2008). To characterize the response of the *AZG2* promoter to auxins, *AZG2_pro_*:GUS and *AZG2_pro_:GFP* activities were studied after application of the exogenous auxin species 1-Naphthaleneacetic acid (NAA). A time course experiment of 3h, 6h, 12h and 24h NAA treatment showed that, after 12 hours of auxin application, GUS staining was enhanced and spread throughout the root, being no longer restricted to LRP environment (Figure 2A). A complementary result was obtained using the *AZG2_pro_:GFP* line (Figures 2B). Moreover, *AZG2* promoter constructs of different lengths, from approximate 2200 to 170bp, were tested for their auxin responsiveness. No differences in expression were observed indicating that there is an auxin-responsive element located within the 170bp upstream of the AZG2 coding secuence start codon. (Supplemental Figures 3B and 3C). As auxin responsive genes are often induced by transcription factors of the auxin response factor (ARF) family (Guilfoyle et al., 1998), we analysed the expression profiles of *AZG2* in the *arf7* and *arf19* mutant background. ARF7 and ARF19 were both linked to lateral root development (Wilmoth et al., 2005). Consistently, *AZG2* (At5g50300) transcription seems to depend on ARF7 or both ARF7 and ARF19 (Okushima et al., 2005). To investigate the relation among ARF7, ARF19 and AZG2 in more detail, *arf7-1, arf19-1* and wild type (Col-0) were transformed with an *AZG2_pro_:RFP* reporter. While no RFP signal was observed in the roots of *arf7-1* mutant (with and without NAA treatment), roots of the *arf19-1* mutant showed the same RFP signal than wild type Col-0 (Figure 2C). Taken together these results demonstrate that the *AZG2* gene is a downstream target of auxin mediated signalling *via* ARF7 transcriptional regulation during lateral root development.

**Figure 3.**
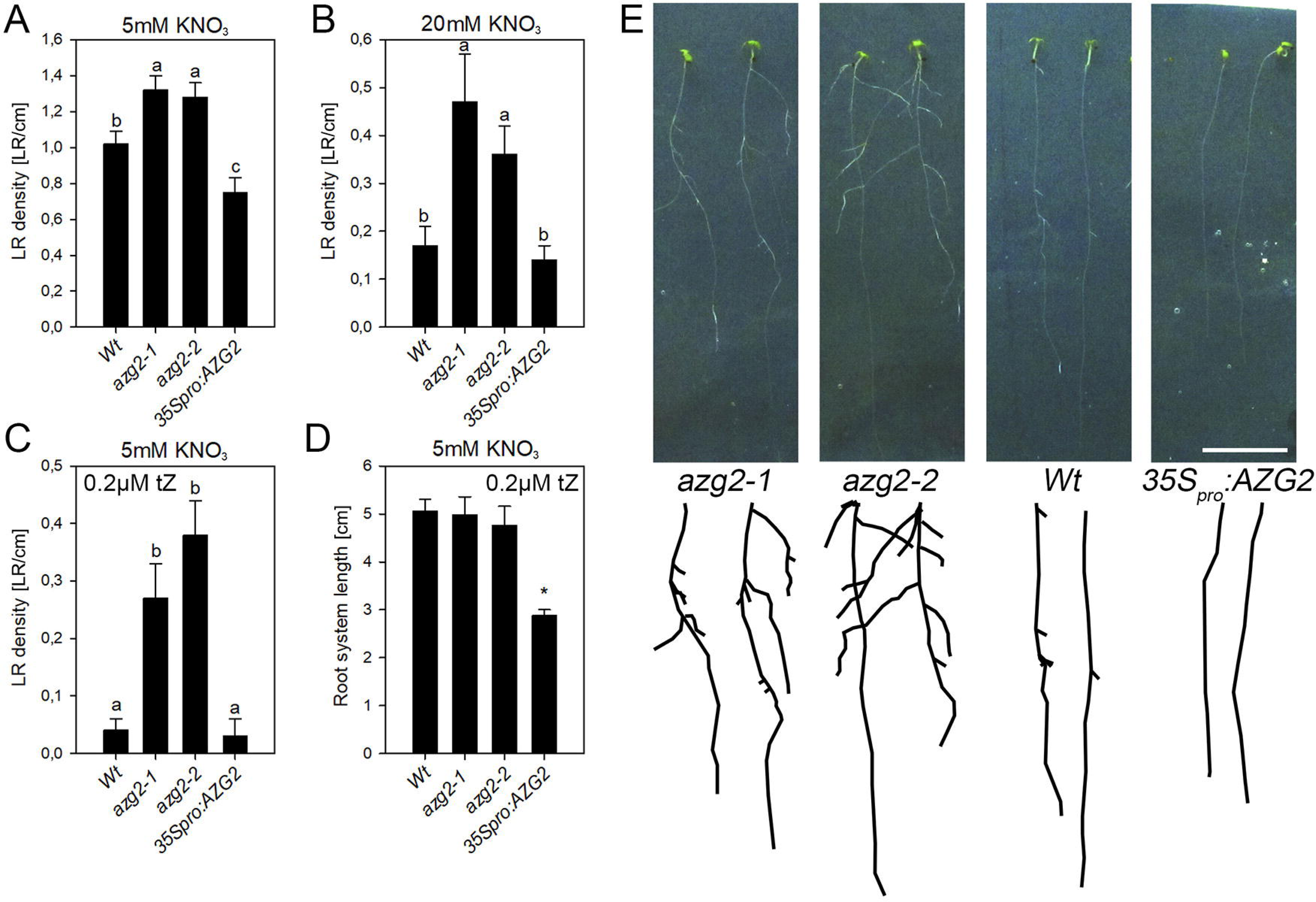
*AtAZG2* mutant lines display altered lateral root densities. (A and **B)** LR density of AZG2 LOF lines and *35s_pro_:AZG2* line in **(A)** 5mM NO^3-^ and **(B)** 20mM NO^3-^ as unique source of nitrogen **(C and D)** LR density **(C)** and total root system length **(D)** of AZG2 LOF lines and *35s_pro_:AZG2* line in 0.5x MS + 0.2 µM tZ. **(E)** Representative image and root skeleton diagram of *Wt, azg2-1, azg2-2* and *35s_pro_:AZG2* plants phenotype growing in presence of 0.2 µM tZ in the media. Scale bar represents 1cm. Values represent the mean ± SEM. Asterisks and letter show significant differences determined by ANOVA test, followed by a Duncan multiple range test (p<0.05 - **(A)** n=105; **(B)** n= 112; **(C)** n=177; **(D)** n= 114).

### AZG2 is part of a lateral root emergence inhibitory pathway

The restricted *AZG2* expression by auxin suggests that the protein may play a role during lateral root development. To address this issue the root phenotype of different *azg2* loss-of-function (LOF) lines (*azg2-1*, *azg2-2*) and a transgenic line ectopically expressing AZG2 (*35Spro:AZG2*) in the wild type background was studied (Supplemental Figures 3A). The *azg2* LOF lines and *35Spro:AZG2* line showed no obvious phenotype under standard growth conditions (0.5x MS). Since it is well known that nitrate availability modulates root architecture (Gruber; 2013; Krouk et al., 2010) and its presence could be masking AZG2 activity, wild type, the LOF and *35Spro:AZG2* line lines were grown in media containing different concentrations of nitrate. Compared to wild type, at 5 mM and 20 mM KNO_3_, *azg2-1* and *azg2-2* plants showed higher lateral root density at 5 mM and 20 mM nitrate (Figures 3A and 3B), whereas the *35S_pro_:AZG2* plants exhibited lower lateral root density at 5mM KNO_3_. No differences were found regarding the primary root length (Supplemental Figure 4C). The opposite phenotypes of the *azg2* LOF mutants and the *35S_pro_:AZG2* line position AZG2 in an inhibitory pathway that controls lateral root emergence.

Since the phytohormone cytokinin is known to have a negative impact on lateral root density (Werner, 2010), the root system phenotype of wild type, the *azg2* mutants and the *35S_pro_:AZG2* line was analysed after growth of the plants on 5 mM KNO_3_-supplemented media containing 200 nM t-zeatin (tZ) in addition. Under this conditions the wild type and *35S_pro_:AZG2* seedlings revealed retarded development, chlorosis and developed nearly none or none lateral roots, respectively (Figures 3C to 3E). Even though the *azg2-1* and *azg2-2* mutants also displayed developmental retardation, they developed a complex root system with several emerged lateral roots. Interestingly, *35S_pro_:AZG2* plants showed a shorter total root system than the wild type and LOF mutant seedlings (Figure 3D). In the *35S_pro_:AZG2* line, but not in wild type or the LOF mutants, *AZG2* is expressed in the root apical meristem (RAM) probably leading to a higher level of CK in the meristem, with a consequent disordered RAM activity. Taken together the cytokinin insensitivity of the *azg2* LOF mutants and *AZG2* expression data and the fact that it is a purine transporter indicate that the AZG2 protein may function in the local distribution of cytokinin (CK).

### AZG2 transports purines with high affinity and CK in plants

To unravel the ability of AZG2 to transport cytokinin, we cloned the *AtAZG1* and *AtAZG2* coding sequences into vectors, which allows the galactose inducible expression of the proteins in yeast (pESC). To verify the functionality of both constructs, the yeast *fcy2* strain, which is deficient in purine transport, was transformed with *pESC-AZG1* or *pESC-AZG2.* Colony growth on medium containing 8-aza-guanine, a toxic analogue of purines was studied. Consistently with the data reported by Mansfield et al. (2009), AZG1 expressing yeast cells showed growth retardation. Surprisingly, no such growth retardation was observed for yeast cells transformed with *pESC-AZG2* (Supplemental Figure 4A). To substantiate the yeast growth phenotypes, we analysed the growth performance of the *Arabidopsis azg2-1* LOF mutants and *35S_pro_:AZG2* lines on growth medium containing either 8-aza-guanine or 8-aza-adenine. As a reference an *azg1-1* mutant and *35S_pro_:AZG1* line were included in this experiment (Supplemental Figure 3B). A developmental retardation and rapid accumulation of anthocyanins were observed for wild type and *azg2-1* mutant seedlings on 8-aza-adenine (Supplemental Figure 4B), whereas *35S_pro_:AZG2* did not show an enhanced toxicity phenotype as it was seen for *35S_pro_:AZG1*. These results suggest that AZG1 and AZG2 have different substrate specificities as it was also suggested by our yeast experiments.

**Figure 4 .**
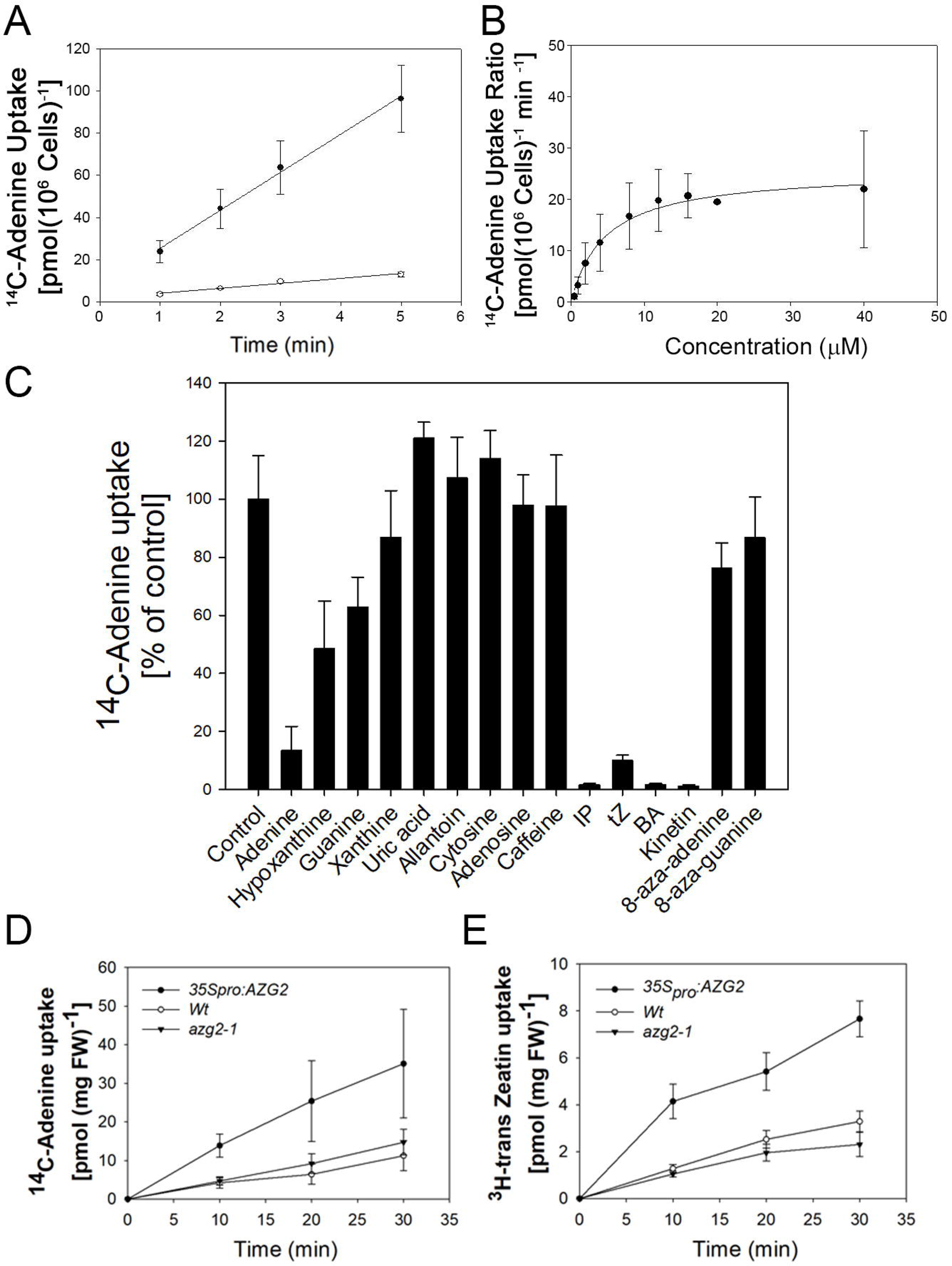
***AtAZG2* transports purine with high affinity in yeasts and is able to take up CK *in planta.*** A yeast strain deficient in adenine transport (*fcy2*) was transformed with AtAZG2 expressed under the control of the GAL 10 promoter in pESC-Ura or with the empty vector. **(A)** Yeast cells transformed with *pESC-AtAzg2* (●) or with the empty vector (○) were assayed for ^14^C-adenine uptake at 20μ substrate concentration and pH 4,5. Radioactivity incorporation after 1, 2, 3 and 5 minutes was measured and expressed in pmol adenine/10^6^ cells. **(B)** AtAZG2-mediated adenine uptake rates (pmol adenine/10^6^ cells/min) were measured at different substrate concentrations. Background uptake rates (empty vector) were subtracted (n= 3) **(C)** Yeast AtAZG2-mediated uptake of ^14^C-adenine (20 μM) was determined in the presence of 10-fold excess (200 μ) of unlabelled compounds. Values are expressed relative to uptake rates obtained without additions (100 %). **(D and E)** Uptake of ^14^C-adenine **(D)** and ^3^H-tZ **(E)** into 14 dag Arabidopsis seedlings. *Wt*(○), *azg2-1* (∇) and *35S_pro_:AZG2* (●) seedlings were incubated with 20μM ^3^H-tZ for the indicated time. Background uptake rates (empty vector) were subtracted. Values represent the mean ± SD of six independent experiments.

In order to check if adenine is a potential substrate for AZG2 in yeast as reported by Mansfield et al. (2009), uptake studies with radiolabeled adenine were carried out. AZG2 mediated linear transport of adenine at least during the initial 5 minutes of incubation and the transport rate was over 10-fold higher compared to the control (empty vector; Figure 4A). The adenine transport rate was saturated by increasing substrate concentration and the deduced Michaelis-Menten kinetic parameters indicated a K_m_ value of 4.75 ± 0.62 μM and a V_max_ value of 25.59 ± 2.05 pmol / 10^6^ cells / min (Figure 4B). The substrate specificity of AZG2 was investigated in ^14^C-adenine competition assays by adding a 10-fold excess of potential unlabelled substrates (Figure 4C). Unlabelled adenine, as was expected, was the purine that strongly inhibited transport, however hypoxanthine and guanine also interfered significantly with the ^14^C-adenine uptake. Cytosine (a substrate for plant PUPs and yeast FCY2), caffeine (transported by PUPs), adenosine (a substrate of ENTs) as well as xanthine, uric acid, uracil and allantoin (substrates for NATs and Ureide Permeases (UPS)) inhibited transport of ^14^C-adenine only slightly. In addition neither 8-aza-guanine nor 8-aza-adenine competed with ^14^C-adenine, which is in line with the previous toxicity experiments (Supplemental Figures 4A and 4B). The published experimental evidence, showing that purine transporters may take a role in CK transport (Burkle et al., 2003; Zürcher et al., 2016) prompted us to evaluate AZG2’s ability to transport ^14^C-adenine in competition with different CK species. Isopentenyl adenine (IP), trans-Zeatin (tZ), benzyladenine (BA) and kinetin inhibited the transport of ^14^C-adenine strongly and even more than the unlabelled adenine itself (Fig 4C). This result indicate that AZG2 may function as a transporter for cytokinins. Nevertheless, the yeast heterologous system has been reported to be elusive in order to analyse CK transport (Ko et al., 2014). Therefore we tested the ^14^C-adenine and ^3^H-tZ uptake into *Arabidopsis* wild type and *azg2-1* as well as in *35S_pro_:AZG2* seedlings. The LOF seedlings showed no significant differences in the the ^14^C-adenine and ^3^H-tZ uptake compared to wild type, while *35S_pro_:AZG2* seedlings exhibited an enhanced uptake capacity for both substances (Figures 4D and 4E). The uptake rate for ^3^H-tZ by the *35S_pro_:AZG2* seedlings was in average 2.5 times higher than that of wild type. Furthermore, the uptake was not saturated even after 30 min of ^3^H-tZ feeding. This result demonstrates that AZG2 is able to transport tZ *in vivo*.

### AZG2 localises to the endoplasmic reticulum and plasma membrane

The elucidation of AZG2’s subcellular localization is a crucial clue to understand its physiological function and to shed light to the discussion about where does the CK signalling begin. In silico analyses showed that AtAZG2 is a membrane protein with around nine to twelve transmembrane domains and has no specific signal peptides for subcellular trafficking (Supplemental Figures 5A to 5C). Transgenic *Arabidopsis* lines were generated that express N- and C-terminal GFP fusions of AZG2 under the control of the Ubiquitin 10 promoter (U10), to trace its subcellular localization. After treating seedlings for short period (10 min) with FM4-64, a partial colocalization of AZG2 and the dye at the plasma membrane of root cells was observed (Figure 5A). In general, the N-terminal and C-terminal GFP fusions of AZG2 showed similar localization patterns within the cells (Figures 5A and 5B). When the cells were additionally treated with BFA, a drug that leads to the accumulation of endomembranes into the so called “BFA compartments”, AZG2 and FM4-64 colocalized in the BFA compartment (Figures 5E to 5G) suggesting that AZG2 gets transported via the TGN on the secretory route to the plasma membrane. Interestingly, beside the PM localization, a GFP signal was also detected inside the cells, especially around the nucleus (Figure 5B, white arrows). Such a protein pattern has been reported as ER localization in roots (Lomin, 2017) and corresponds with the signal for ER-targeted GFP (Figure 5 up-right panel; Ottenschläger et al., 2003; Müller and Sheen, 2008). To verify that the observed pattern correspond to the ER, a transgenic *Arabidopsis* line was analysed that express AZG2-GFP under the control of the 35S promoter. This overexpression line exhibited a GFP signal in leaf epidermal cells that is characteristic for an ER pattern (Figure 5C).

**Figure 5.**
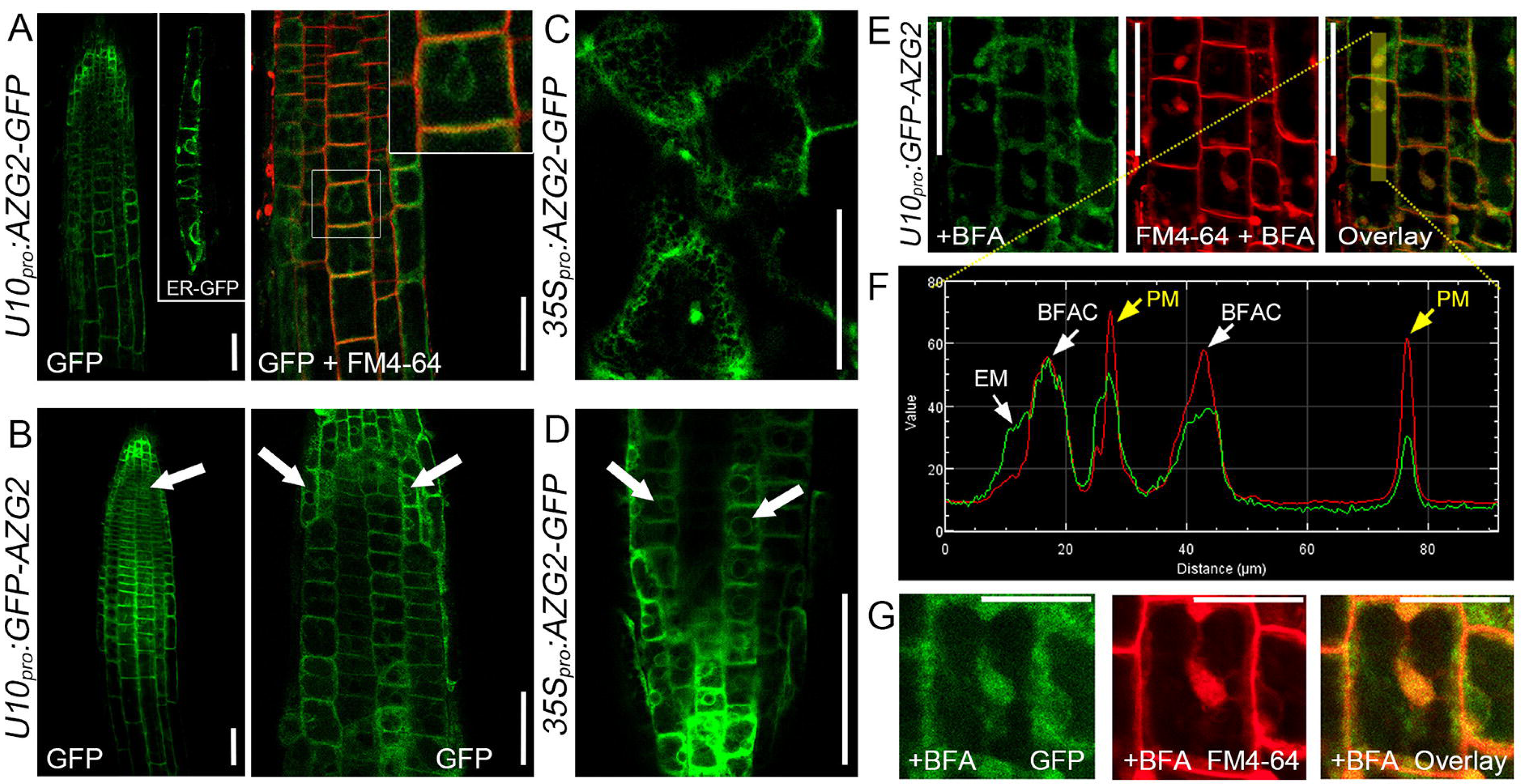
***AtAZG2* is localised in plasma membrane and in endoplasmic reticulum**. **(A)** Carboxyl and **(B)** amino terminal fusion of AZG2 with GFP expressed under the control of Ubiquitin-10 promoter in the main root tip of 7 dag seedlings. ER-GFP corresponds to a GFP tagged for ER retention under the control of *TCSn_pro_*. White arrows shows perinuclear signal. FM4-64 short treatment label plasma membrane. **(C and D)** Overexpression of AZG2-GFP using 35S promoter showing AZG2 signal in **(C)** leaf epidermal cell and **(D)** in root tip. **(E)** Seedlings of *pU10:GFP-Azg2* after 1 hour BFA 1mM treatment followed by a 10 minutes incubation with FM4-64. **(F)** Fluorescence quantification in GFP (green) and FM4-64 (red) channels within the highlighted region in yellow **(E)** (PM: plasma membrane; BFAC: Brefeldin-A compartment; EM: endomembrane). **(G)** Detail of a cell in the root elongation zone treated with BFA. Scale bars represents (A-D) 50 µm and 25 µm (G).

It has been argued that gene expression under 35S promoter results in experimental artifacts by leading protein stuck in the ER. However, the root signal pattern (Figure 5D) is coincident with that observed in the lower expression lines *U10:AZG2-GFP* and *U10:GFP-AZG2* (Figure 5A and 5B) suggesting that the observed pattern is not product of an overexpression artifact.

### The absence of AZG2 compromises CK signalling in root

In order to gain deeper insights into AZG2 function in CK signalling, the activity of the CK signalling reporter *TCSn_pro_:GFP* (Zürcher et al., 2013) was studied in wild type and *azg2-1* genetic background. As shown in Figure 6A the *TCSn_pro_:GFP* reporter gene was active in both genetic backgrounds under control conditions but the GFP signal was lower in the roots of the *azg2-1* mutant than in the roots of wild type. After 24 h of tZ exposure, the GFP signal increased in both genotypes but was lower in the roots of the *azg2-1* mutant compared to wild type. This differential GFP intensity can be specifically noticed in the cells covering the primordia, where AZG2 is normally expressed (Figure 6A; white arrows). In order to quantify *TCSn_pro_* activity, the GFP signal of two independent events of *TCSn_pro_:GFP* in the *azg2-1* mutant background was analysed and compared to the signal in wild type. The GFP signal was significantly lower in the *azg2-1* mutant roots compared to the signal of wild type roots (Figures 6B and 6C). This results reinforce the hypothesis that AZG2 functions as a CK transporter involved in root hormone signalling.

**Figure 6.**
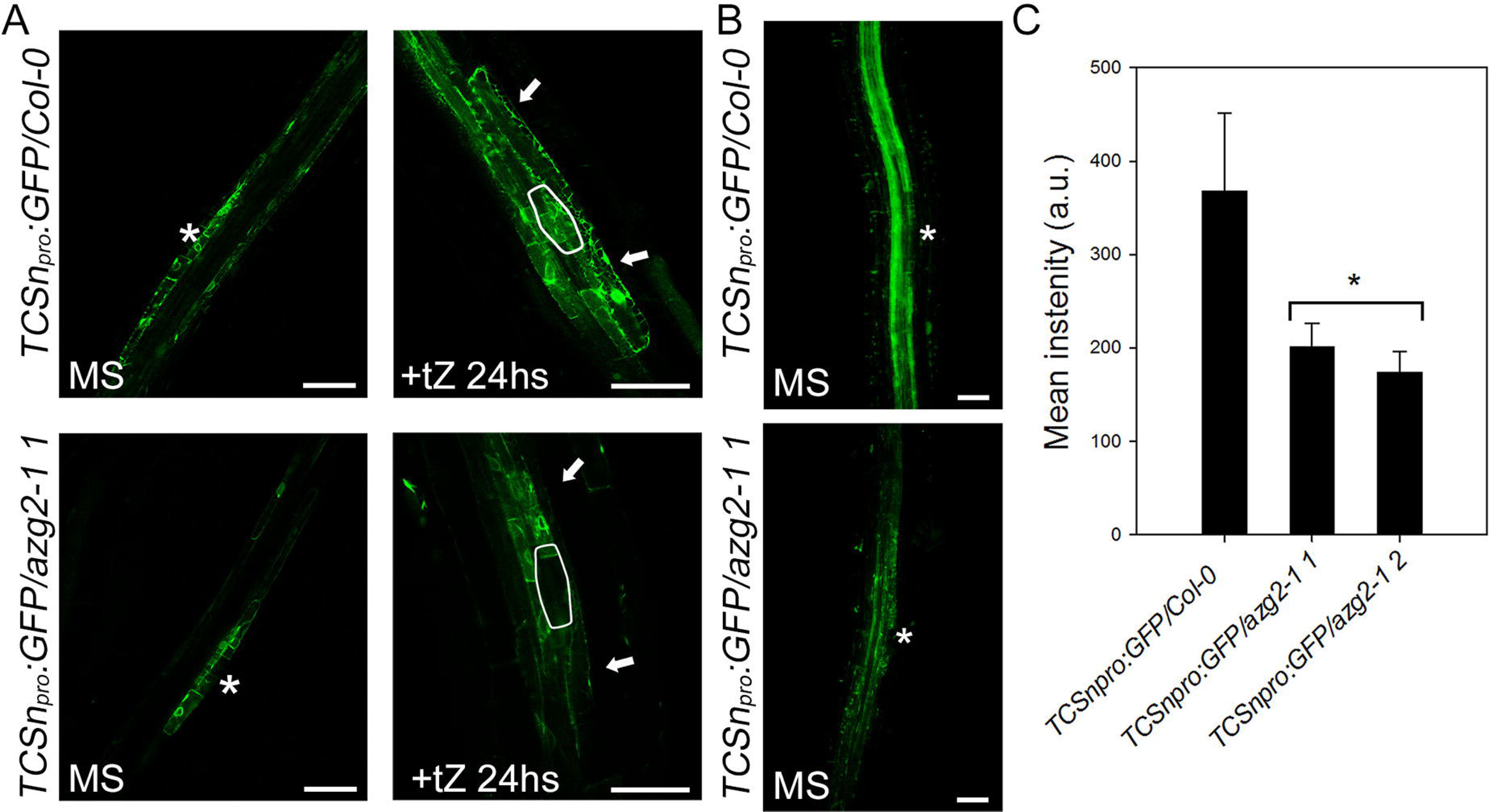
CK signalling is lower in AZG2 KOs. (A) Expression of *TCSn_pro_:GFP* in Col-0 and *azg2-1* background with and without 24 hours 0,2uM tZ treatment. White line and asterisks delimit the LR primordia. **(B)** Representative z-projection used for fluorescence quantification of Col-0 and *azg2-1* expressing *TCSn_pro_::GFP* grown in 0.5x MS. **(C)** Fluorescence quantification of GFP in roots z-projections. Six dag seedlings of Col-0 and two independent insertion lines of *azg2-1* grown in 0.5x MS were imaged. Values represent the mean ± SEM. Asterisk shows significant differences determined by ANOVA test, followed by a Duncan multiple range test (p< 0.05; n= 27). Scale bars represent 50uM.

### AZG2 facilitates CK diffusion through membranes

As demonstrated previously and by our experiments using the yeast *fcy2* mutant, AZG1 transports 8-aza-guanine and 8-aza-adenine with higher efficiency than AZG2, pointing to the possibility that they have different transport properties.

We therefore measured the uptake of ^14^C-adenine into yeast *fcy2* cells at different extracellular pH values. AZG1 showed maximal uptake activity at pH 4 which descreased with increasing pH the (Figure 7A). In contrast, AZG2-dependent adenine uptake increases with increasing extracellular pH and reached saturation at pH 5 (Figure 7B). In the absence of an energy source in the media (-Galactose), in the presence of the protonophore carbonyl cyanide m-chlorophenyl hydrazone (CCCP) or the ATPase inhibitor diethylstilbestrol (DES), the adenine uptake decreased independent whether the yeast cells express AZG1 or AZG2 (Figure 7C). Nevertheless the dependence of the adenine uptake on an energy source and the existence of an electrochemical proton gradient over the plasma membrane was less pronounced for AZG2 than AZG1 (Figure 7C). These data suggests that AZG1 mediates an H^+^-driven adenine transport similarly to AZGA in *Emerichella*. On the contrary, AZG2 displays a dynamic of facilitated transport, being not dependent as much on the electrochemical proton gradient or an energy source. Finally, to better understand the transport features of AZG1 and AZG2 even more, yeast cell expressing AZG1 or AZG2 were fed for 10 min with radiolabelled hypoxanthine, an additional purine transported by AZG2 with high affinity (K_m_=11,16 ± 1,34 µM; Supplemental Figure 6). After several washings, the remaining radioactivity inside the cells was measured after 0 (100%), 3, 5 10 and 20 minutes incubation in pH 4.5 for AZG1 and pH 7 for AZG2. Results showed that AZG2 lost the double compared to AZG1, with a drop of around 40% of the label (Figure 7D). All these results together suggest that while AZG1 seems to rely on proton coupled uptake, AZG2 could behave as diffusion facilitator.

**Figure 7.**
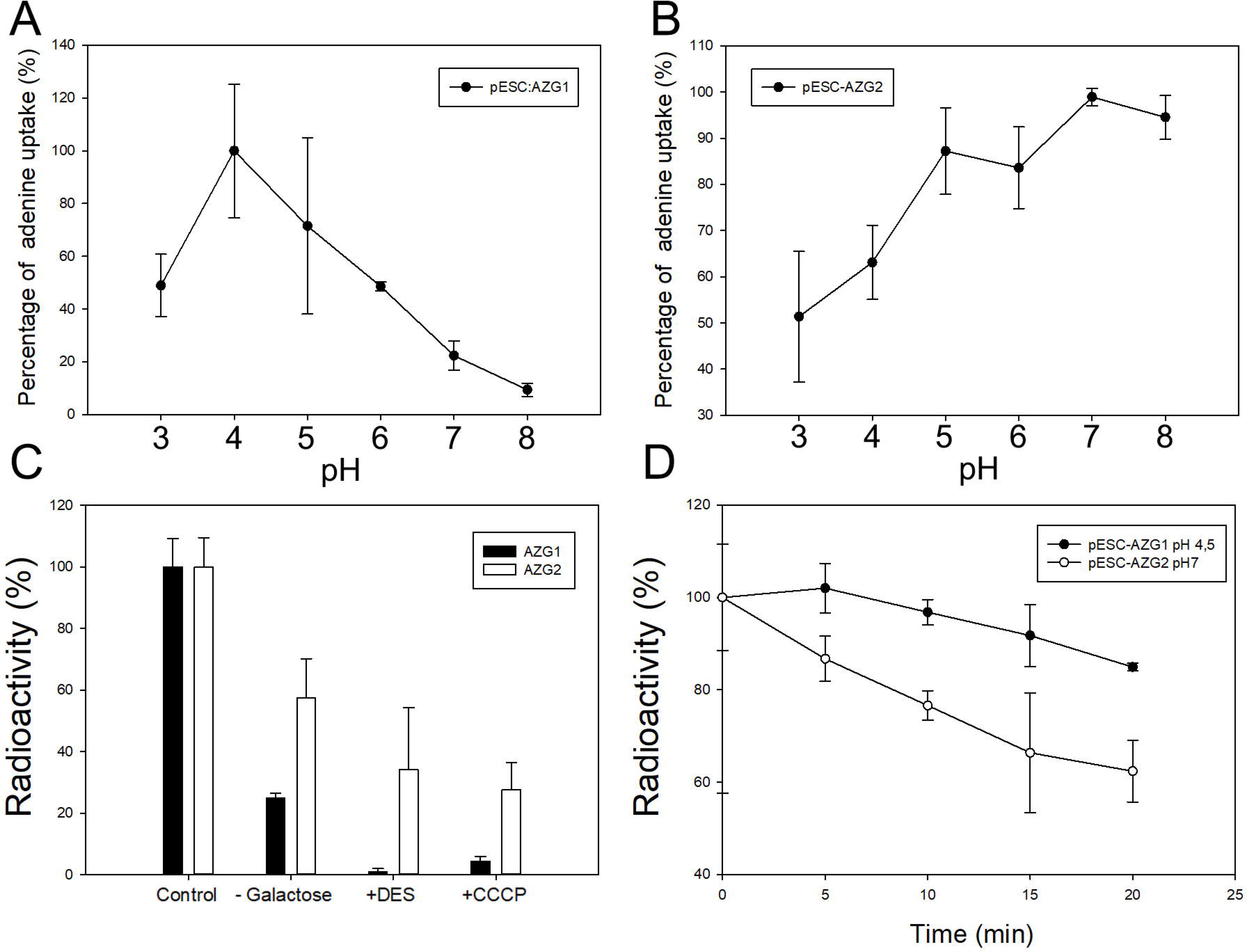
*AtAZG2* acts as a transport facilitator and its activity is independent of an energy source. (A and B) ^14^C-adenine uptake measured under different pHs in yeast (*fcy2*) carrying **(A)** *pESC:AZG1* and **(B)** *pESC:AZG2*. Percentage is relative to the maximum uptake value. **(C)** AZG1 (black) and AZG2 (grey) mediated ^14^C-adenine uptake rate measured under standard conditions, excluding galactose as energy source from the preincubation medium (-Gal), or in the presence of 100 μM CCCP and DES. Percentage of adenine uptake is relative to the control. **(D)** Yeast cell expression AZG1 or AZG2 fed for 10 min with ^3^H-hypoxanthine. Then cell were washed and incubated in cold buffer for 5, 10, 15 and 20 minutes to evaluate ^3^H-hypoxanthine efflux at pH 4,5 (● AZG1) and pH 7 (○ AZG2). Percentage of hypoxanthine uptake is relative t_0_ value. Values represent the mean ± SD of 3 independent experiments.

## DISCUSSION

Root system architecture is determined by root branching that is highly regulated by hormones, mainly auxin and CK. During the last decade, hormone transport became a key factor in the understanding of plant hormone homeostasis. To date two CK transporters, ABCG14 and PUP14, were described to be involved in the modulation of plant development. ABCG14 is expressed in the endodermis and stelar cells, acting as an efflux carrier for long distance CK transport (Ko et al., 2014; Zhang et al., 2014). On the other hand, PUP14 is expressed throughout the plant and has been proposed as a transporter which modulates the efficiency of CK signalling at plasma membrane level (Zürcher et al., 2016). In this study, we introduce AZG2 as a putative CK transporter, which has a very restricted expression domain in few cells covering the lateral root primordia (Figure 1). As OLT plays a key regulatory role on primordia development (Peret et al., 2012; Lucas et al., 2013), AZG2 arises as a suitable candidate for the control of local CK gradients in the lateral root environment.

### AZG2 is a candidate for the supply of CK to the cytokinin receptors located in the ER

Purine transporters such as PUP1 and PUP14 are able to transport not only adenine, but also CK (Burkle et al., 2003; Zürcher et al., 2016). Mansfield (2009) showed that AZG transporters from *Arabidopsis* are able to take up adenine and guanine in yeast and *Arabidopsis* seedlings. In this work, we demonstrated that AZG2 additionally transports CK in yeast and *Arabidopsis* with high efficiency. When AZG2 CK accumulation was determined in *Arabidopsis* seedlings, no differences were found between the *azg2* LOF mutants and wild type seedlings. This is probably due to the spatially very restricted activity of the AZG2, as the minor addition of CK to a few cells cannot be detected against the background of the entire seedling. In contrast, the *35S_pro_:AZG2* line took up significantly more labelled tZ than wild type showing that AZG2 is able to transport CK *in vivo*. To understand a potential role of AZG2 in the CK signalling pathway, it is necessary to determine which cell compartments become connected by its activity. Therefore, the subcellular localization was traced in transgenic *Arabidopsis* plants by using different GFP fusions of AZG2. All fusion proteins located to both the plasma membrane and the ER. These observations arose the questions if AZG2 is functional at the conditions given at both membranes sides. Studies in yeast indicated that AZG2 has maximal transport activity in media with pHs between 5 to 8, contrasting with AZG1 which shows clear transport dependence on an electrochemical proton gradient (Fig. 7A-B). Particularly, the capacity of AZG2 to efficiently transport substrates even at slight alkaline pHs fits very well with a functional transporter at the ER. Additionally, the export studies in yeast suggest that AZG2 is able to move their substrates in both directions (Fig. 7D). Thus, AZG2 may function *in vivo* as a diffusion facilitator, whereas AZG1 may perform secondary active transport. Taken together, these results support the presence of functional AZG2 in both PM and ER of cells expressing the transporter. The latter observation is relevant regarding the ongoing discussion about the subcellular localization of the functional CK receptors (Caesar et al., 2011; Romanov et al., 2018). Here, AZG2 KOs partial insensitiveness to exogenous CK (Figure 3C) becomes a key to unravel the transporter biological relevance. Although *azg2* LOF mutants still can take up exogenous CK, probably by other transporter such as PUP14 (Zücher et al., 2016), they display a defect in CK-controlled lateral root emergence. Against the background of the spatially restricted expression, the loss of AZG2 function could interfere with the CK supply from the plasma membrane to the ER lumen, where the ligand-binding domain of the CK receptors is exposed (Figure 8A). The interruption of the CK supply, then leads to CK insensitivity in the OLT cells and eventually to the impaired lateral root emergence observed for the *azg2* LOF mutants.

**Figure 8.**
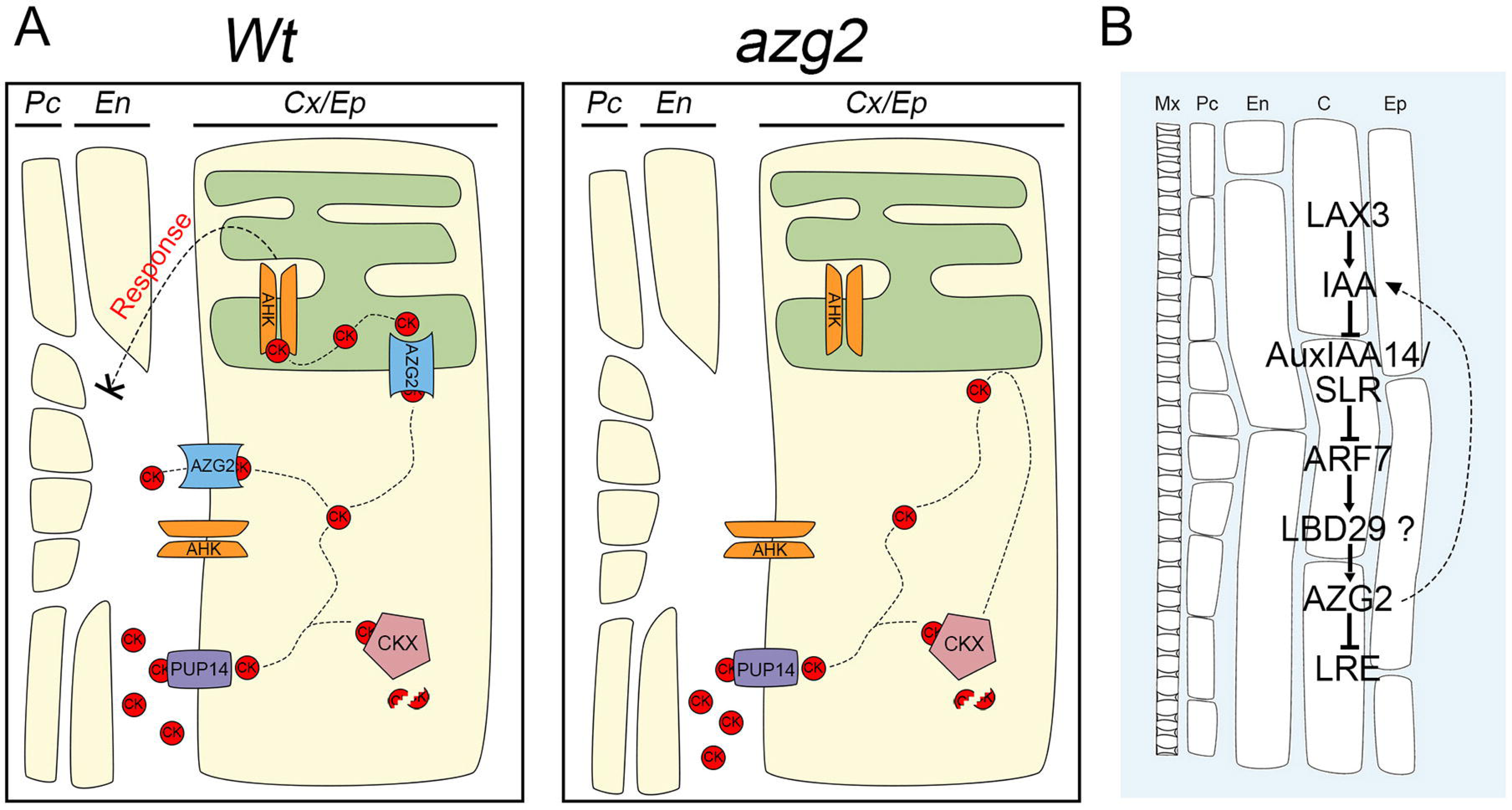
Theoretical signalling model (A) In *Wt* background genotype, PUP14 transporter could take up CK with high affinity preventing CK to reach PM AHKs. This function can be also lead by AZG2 transporters localized in plasma membrane of OLT cells. Once in the cytosol, CK can alternatively be degraded by cytosolic CKX enzymes, or be transported into the ER lumen via AZG2 to reach intracelular AHKs triggering the CK signalling cascade which regulates LR emergence. On the other hand, in *AZG2* KO background, the absence of AZG2 results in an CK accumulation in the cytosol as a sink, being unable to reach AHKs sensitive domine. This results in an altered regulation of LR emergence. **(B)** AZG2 is part of a Auxin/CK signalling network. This Auxin/CK signalling loop begins with the auxin import by LAX3 in OLT releasing ARF7. Then, ARF7 enhanced LDB29 (or other LBD) expression, which induces AZG2 expression. The activity of AZG2 transporter through still unknown mechanisms, acts as a LR emergence (LRE) inhibitor. Besides the change in CK levels might have impact in the auxin homeostasis. (Mx: Metaxylem; Pc: Pericycle; En: Endodermis; Cx: Cortex; Ep: Epidermis).

### AZG2 is part of a lateral root emergence inhibitory pathway

We found that the promoter region 170bp upstream of the start codon were sufficient for the auxin inducibility of the *AZG2* gene. AZG2 expression dependence on auxin was previously mentioned by Okushima et al. (2005) who included *AZG2* among the genes induced by ARF7, but not ARF19. This was confirmed in this work by studying *AZG2_pro_:RFP* expression in *arf7-1* and *arf19-1* mutant background. However, within these 170bp, no canonical auxin response element (*AuxRE*; Ballas et al., 1993) is present, suggesting that either another but ARF7-dependent transcription factor or a non-canonical *AuxRE* might be involved in the *AZG2* regulation. Interestingly, 165bp upstream of the start codon a G-box sequence (5’-CACGTG-3’) is present in the *AZG2* promoter. This sequence has been reported as a target sequence for the LBD29 transcription factor (Xu et al., 2017), which acts downstream of ARF7 (Okushima et al., 2007). Nevertheless, other LBDs cannot be excluded in *AZG2* gene regulation, since they modulate expression of many genes involved in root architecture (Feng et al., 2012; Lee et al., 2015; Porco et al., 2016). Indeed, LBDs induce the expression of several genes in the OLT such as those encoding the cell wall remodelling enzymes EXP14 and EXP17, which are required for lateral root primordia emergence (Lee and Kim, 2013; Lee et al., 2013). Here, it is relevant to highlight that the OLT and the micropylar endosperm, both tissues where AZG2 expressed (Supplemental Figures 2E to 2G), have a high cell wall remodelling activity. Although it is known that CK regulate several genes involved in cell wall remodelling, it physiological relevance during development has not been studied yet (Brenner et al., 2012). This could be one of the processes downstream of AZG2 that influence lateral root emergence. In concordance, the root system of *azg2* LOF lines developed more lateral roots than that of the wild type and *35S_pro_:AZG2* plants. Whether this inhibitory pathway is involved in the emergence or other events in lateral root development is motive of present investigations.

Both, the auxin-dependent induction and the CK transport activity of AZG2 suggest that this transporter is part of the auxin/cytokinin hormonal crosstalk. The direct relevance of AZG2 function in CK signalling is evident as *TCSn_pro_:GFP/azg2-1* roots showed less CK signalling activity around LRPs than wild type. Local activation of CK *via* biosynthesis induced by auxins has been reported by De Rybel et al. (2014). Within this auxin/CK loop, LAX3 (<u>L</u>ike <u>A</u>U<u>X</u>1 <u>3</u>), which is expressed in the OLT, may transports auxins into the OLT cells that release ARF7 from AUX/IAA14 inhibition (Fukaki and Tasaka, 2009). Then, ARF7-upregulated LBD transcription factors induce the expression of the CK transporter AZG2 (Figure 8B). Thus, AZG2 would be part of a lateral root emergence inhibitory pathway, reinforcing the antagonistic role of auxins and CK. Despite the fact that downstream events of AZG2 transport activity are unknown, regulation of cell wall remodelling is a promissory research aim to gain insight into AZG2 activity during lateral root emergence. This could help to explain how AZG2 along with other positive and negative regulators comprise a fine tuned lateral root emergence regulation mechanism.

## METHODS

### Plant material

*Arabidopsis thaliana* accession Columbia (Col-0) was used as wild-type. AZG1 and AZG2 knock-out lines were obtained from Nottingham Arabidopsis Stock Center (http://arabidopsis.info/). Two independent T-DNA insertion lines of AZG1, SAIL_114E03 (*azg1-1*) and GK-681A06 (*azg1-2),* and two independent T-DNA insertion lines of AZG2, SALK 000904 (*azg2-1*) and Sail 658 G02 (*azg2-2*), were genotyped. The presence of T-DNA insertion was checked by PCR using the following primers couples; for *azg1.1* Azg1-R and SAIL-LB3 (5’-TAGCATCTGAATTTCATAACCAATCTCGATACAC-3’); for *azg1.2* Azg1-R and GABI-Kat-LB (5’-ATATTGACCATCATACTCATTGC-3’); for *azg2.1* SALK-LBb1.3 (5’-ATTTTGCCGATTTCGGAAC-3’) and Azg2-R, for azg2.2 Azg2-F and SAIL-LB1 (5’-GCCTTTTCAGAAATGGATAAATAGCCTTGCTTCC-3’); (Supplemental Figure 1A and 1B). The presence of the Azg1 and Azg2 wild type alleles -by PCR- and the absence of mRNA -by RT-PCR- was checked with Azg1-F (5’-CACCATGGAGCAACAGCAACAAACA-3’) and Azg1-R (5’-ATTGGGGAGAA GAAGGTTTGGTGAAAT-3’); and GW-Azg2-F (5’-GGGGACAAGTTTGTACAAAAAAGCAGGCTTAATGACATCCTCCTCCTGCAT TTGC-3’) and GW-Azg2-R (5’-GGGGACCACTTTGTACAAGAAAGCTGGGTCTCAAACGATCTCGACGGTAGCGAC-3’) primers respectively (Supplemental Figure 1C and 1D). AZG1 and AZG2 overexpression lines were obtained from Dr. Frommer laboratory. The *arf7-1* and *arf19-1* alleles were used in this study as knock-out mutants of ARF7 and ARF19. Both knock-out lines *arf7* (N24607) and *arf19* (N24617) were obtained from NASC.

### Growth conditions

Plants were cultivated under short day photoperiod conditions (8hs light /16hs dark illumination regime) at 23°C for root phenotype analysis and long day photoperiod for the rest (16hs light /8hs dark). For in vitro cultures, seed were superficially sterilized for 10 minutes with ethanol 70% + 0.05% Triton X-100, followed by two washings with ethanol 96% of 10 minutes and sowed immediately after drying. To allow gas exchange and prevent fast desiccation, plates were sealed with Micropore™ tape. Imbibed seeds were stratified at 4°C for at least 48 h. Culture media, specific nutritional conditions are describe below for each experiment.

### Expression in yeast and uptake assays

AtAzg1 coding region was amplified using the following primers (5’-ATACCCGGGGAATTCATGGAGCAACAGCAACAACAAC-3’) and (5’-GCGGCCGCGTTTTTCTTTGATTTGATTTTAC-3’) with the EcoRI and NotI sites. The excised fragment was introduced in the MCS 2 (EcoRI/NotI) of the yeast expression vector pESC-URA (Stratagene), cloned in the E. coli strain DH5-α and sequenced. Besides, AtAzg2 coding region was amplified using (5’-ATACCCGGGACTAGTATGACATCCTCCTCCTGCATTTGC-3’) and (5’-ATATCTAGATCAAACGATCTCGACGGTAGCGAC-3’) in cloned into pOO2 and sequenced. Then, Azg2 CDS was subcloned into pESC-URA using XhoI and SmaI restriction sites. Both constructs were introduced into the yeast mutant strain MG887-1 (Mat a, fcy2, ura3; Gillissen et al., 2000) as previously described (Gietz and Schiestl, 1994). Transformed yeast cells were grown under selective conditions in minimal medium containing CSM-URA and glucose as carbon source. For transport studies cells were grown in medium with galactose as carbon source and harvested at an OD600 of 0.8 by centrifugation for 5 min at 4500 rpm. Cells were washed and resuspended in double distilled water to a calculated final OD600 of 20. 165 μl of cells were mixed with 33 μl of 1M potassium phosphate, pH 4.0, 33 μl of 1 M galactose, and 66 μl of double-distilled water and preincubated for 5 min at 30° C. To start the reaction, 33 μl of 200 μM of the respective radiolabeled substrate was added. Samples of 100 μl were removed after selected times, transferred to 4 ml of ice-cold water, filtered on fiberglass filters, and washed with 8 ml of water. Radioactivity incorporated in cells was determined by liquid scintillation spectrometry (Beckman). Transport measurements were repeated independently at least 3 times. Radioactive substrates were: ([8-14C]-adenine (10.8 GBq/mmol; Amersham Biosciences UK Limited); [8-3H]-hypoxanthine (925 GBq/mmol; GE Healthcare UK Limited); trans-[2-3H]-zeatin (592 GBq/mmol; Institute of Experimental Botany of the Czech Academy of Sciences, Isotope Laboratory, Praha). Radiochemical purity of trans-[2-3H]-zeatin was determined before use basically as described previously (Burkle et al., 2003).

### Uptake assays with *Arabidopsis thaliana* seedlings

The uptake of radiolabeled substrates into Arabidopsis plants was determined using a modification of a previously described method (Sherson et al., 2000). Briefly, Arabidopsis seeds of the wild-type, T-DNA insertion lines *azg1-1, azg1-2, azg2-1* and *azg2-2* and overexpressing lines *35S_pro_:AZG1* and *35S_pro_:AZG2* were sterilized and plated on MS-agar plates. After two weeks, seedlings were collected for a minimum weight of 5 mg and a minimum number of four plants for each reaction tube. The plants were vacuum infiltrated in 450 μl MS-medium (pH adjusted to 4.5) at 300 mbar for 3 min followed by incubation in MS media for 30 min at RT for equilibration. 50 μl of the respective radiolabeled substrate (25μM final concentration) were added and incubated for 10, 20 or 30 min. The plants were washed five times with 4 ml ice-cold MS-medium for one minute each. Finally, samples were extracted in 80 % ethanol and radioactivity determined by liquid scintillation counting. Transport measurements were repeated independently and represent the means of at least six experiments. Radioactive substrates were: ([8-14C]-adenine (10.8 GBq/mmol; Amersham Biosciences UK Limited); trans-[2-3H]-zeatin (592 GBq/mmol; Institute of Experimental Botany of the Czech Academy of Sciences, Isotope Laboratory, Prague).

### Constructs for Azg2 tissue and subcellular localization

To address Azg2 expression in plant organs and tissues, Azg2 1.5kb promoter was cloned to drive GUS reporter (pGPTV-bar). To study the active length of AZG2_pro_, different lengths of AZG2 promoter (−2149, −1388, −940, −169bp upstream from start codon) were chosen to drive GFP expression. The four amplicons were cloned into pENTR™/D-TOPO® (Invitrogen) via BP reaction and the subsequent LR reaction into pPGG and pPGR gateway vectors with the mGFP5 and RFP marker respectively. As well, to study subcellular localization AZG2 fusions with GFP or RFP were generated. *AtAzg2* CDS was clone into *pDONR207* entry vector and subcloned into *pUBC-GW-GFP-DEST*, *pUBC-GFP-GW-DEST* to express AZG2 constitutively with a reporter N- and C-terminal fusion. For overexpression with CMV35S promoter *AtAzg2* CDS was cloned into pDONR221 and subclone in *pFRETcg-2in1-CC* generation a C-terminal fusion with GFP. All constructs were sequenced to discard mutation. For all transformations plasmids were introduced in *Agrobacterium tumefaciens* strain C58/ATCC33970. Five to six weeks old plants were transformed using floral dip method as described Clough and Bent (1998). Mutant plants were selected with the correspondent antibiotic and genotyped.

### GUS staining

Histochemical analysis of the GUS reporter enzyme was performed as described Martin et al. (1992), with minor modifications. For extensive expression description 19 days after germination (dag) seedlings grown on Petri dishes and reproductive organs of adult plants growing in soil were stained. AZG2_pro_:GUS plants were incubated in staining solution overnight followed by 3 subsequent incubations with 80% ethanol. Stained material was mounted and analysed and photographed under DIC microscopy with Olympus BX61 microscope. Olympus DP71 camera was used for image acquisition.

### Semi-quantitative RT-PCR analysis

To evaluate the expression of the AtAzg2, total RNA from different organs of *Arabidopsis thaliana* were isolated from 100 mg tissue using the TRIzol reagent (Gibco-BRL). RNA was converted into first strand cDNA using the SuperScriptII Reverse Transcriptase (Invitrogen). PCR reactions were conducted in a final volume of 10 ul using 1 ul of the transcribed product and Taq DNA polymerase (Quiagen). The pairs of primers used were Azg2-SmaI-F and Azg2-XhoI-R. Amplification conditions were as follows: 3 min denaturation at 94°C; 35 cycles at 94°C for 30 s, 53°C for 40 s and 72°C for 30 s, followed by 5 min at 72°C. As control, the Actin2 gene was amplified by 28 cycles and the following primers were used: Actin2-F (5’-TGTACGCCAGTGGTCCTACAACC-3’) and Actin2-R (5’-GAAGCAAGAATGGAACCACCG-3’).

### Hormone treatments

For auxin induction 10 dag seedlings of *AZG2_pro_:GUS* were transfer to liquid 0.5x MS pH5.8 media with 1µM naphthalene acetic acid (NAA - PhytoTechnology) for 3, 6, 12 and 24 hours. For seedlings carrying GFP or RFR reporter, 10 dag seedlings of *AZG2_pro170_:GFP*, *AZG2_pro1000_:GFP*, *AZG2_pro1400_:GFP*, *AZG2_pro2200_:GFP*, and *AZG2_pro_:RFP* were incubated for 12hs and immediately observed under Nikon Eclipse Ti confocal microscope. For CK 24hs treatments, plant were transfer from 0.5x MS pH5.8 plates to liquid 0.5x MS pH5.8 media with no trans-Zeatin (MS control) and 0.2µM trans-Zeatin (Duchefa). For CK long term treatment, plants were grown under 0.5x MS + KNO3 5mM + 0,2µM trans-Zeatin.

### AZG2 subcellular localization and *TCSn:GFP* signal quantification

Six dag seedlings of different lines growing in 0.5x MS under short photoperiod were analysed with Nikon Eclipse Ti confocal microscope. Scan was performed with 20x and 60x oil immersion lens and with a 1024×1024 pixel resolution. For *U10_pro_:AZG2-GFP* and *U10_pro_:GFP-AZG2* samples were scanned with a 488 nm laser and emission detected with a 515/30 filter. For BFA treatments *U10_pro_:GFP-AZG2* seedlings were incubated for incubated for 1 hour with 10μM FM4-64 + 50μM Brefeldin A. Shot FM4-64 incubation of 10 minutes were performed in order to stain the plasma membrane.

The characterized line *TCSn_pro_:GFP/Col-0* and the *TCSn_pro_:GFP* plasmid construct were kindly provide by Dr. Müller laboratory. Knock-out line *azg2-1* was transformed as described above with *TCSn:GFP* construct via floral dip and T3 plants were selected in homozygosis. Two independent insertional lines were selected and analized to discard insertional variation of the expression. Nine dag plants of *TCSn_pro_:GFP/Col-0, TCSn_pro_:GFP/azg2-4 1* and *TCSn_pro_:GFP/azg2-4 2* were observed under Nikon Eclipse Ti microscope. To quantify fluorescence intensity, LPRs of seedlings were imaged with a 20x objective and 1024×1024 pixel resolution. Each sample image represents a z-stack of 12 steps throughout the root cylinder centred in the LRP. Stacks were z-projected (average intensity) and mean intensity was quantified. Images were processed and analysed using ImageJ-FIJI.

### Plants phenotype description

For toxicity essay *Arabidopsis thaliana* seedlings were first sown on MS medium containing 1% sucrose (w/v) for 10 days and then alternatively transferred to greenhouse conditions or to plates containing MS supplemented with 50 μM 8-aza-adenine or 8-aza-guanine or 0.2 μM tZ.

For root architecture studies plants were grown in 0.5x MS without nitrogen (PhytoTechnology) + 5 mM and 20 mM KNO_3_ for 19 days in 120mm square plates. Seed of all lines belong to the same bath to avoid any unwanted variation, as they also were grown side by side under uniform illumination. Plates were photographed with Canon PowerShot SX150IS camera, and roots were measure with ImageJ. For phenotype analysis under exogenous CK exposition the same procedure was carried out and plants were photographed at 19 dag.

---

**Supplemental Figure 1. (A and B)** Genotyping of **(A)** *azg2* and **(B)** *azg1* alleles by PCR using combinations of gene specific primers (g) or T-DNA specific primers (i) (see Methods). **(C and D)** RT-PCR analysis on mRNA isolated from **(C)** *Azg1* and **(D)** *Azg2* knockout lines and controls. As control the specific genomic fragment was amplified from the *Wt* background.**(E)** Expression of AtAZG2 in different tissues and developmental stages obtained from EFP Browser (http://bar.utoronto.ca/efp/cgi-bin/efpWeb.cgi; Winter, 2007)

**Supplemental Figure 2. (A to D)** *AZG2_pro_:GUS* plants **(A and B)** different states of flower development, **(C)** pollen and **(D)** immature seeds do not show reporter expression. **(E to G)** *AZG2_pro_:GUS* germinating seeds stained, showing signal under seed tegument after **(E and F)** 8 and **(G)** 24 hours light exposition. **(F)** Seed section showing GUS signal under seed tegument (arrow).

**Supplemental Figure 3. (A)** Induction of AtAZG2 expression after a 3 h auxin (IAA) treatment (Winter, 2007). **(B and C)** AZG2 expression in OLT. **(A)** *AZG2_pro_:GFP* expression in cortical cell surrounding LR primordia. **(B)** Different promoter lengths (170, 900, 1400 and 2200bp) shows expression in LR OLT. Scale bars represents 100µm.

**Supplemental Figure 4. (A)** Toxicity essay in yeast transformed with the pESC empty vector, pESC-AZG1 and pESC-AZG2 exposed to increasing concentration of 8-aza-guanine. **(B)** AZG1 and AZG2 KOs and OE lines response to 50µM 8-aza-adenine and 8-aza-adenine in the growing media. **(C)** Main root length of AZG2 mutant lines in MS 0.5x with 5mM KNO_3_ as single nitrogen source.

**Supplemental Figure 5. (A)** Number of transmembrane domains of AtAZG2 transporter estimated by TMHMM (Krogh, 2001). **(B)** Estimation of transmembrane domains by different algorithms available at ARAMEMNON (http://aramemnon.uni-koeln.de). **(C)** Signal peptide prediction using SignalP-4.1 (Petersen, 2011) suggests that AtAZG2 lacks of a signal peptide.

**Supplemental Figure 6.** AtAZG2-mediated hypoxanthine uptake rates (pmol adenine/10^6^ cells/min) were measured at different substrate concentrations. Background uptake rates (empty vector) were subtracted (n= 3).

